# Deciphering *Staphylococcus aureus-*host dynamics using dual activity-based protein profiling of ATP-interacting proteins

**DOI:** 10.1101/2024.02.05.578939

**Authors:** Stephen Dela Ahator, Kristin Hegstad, Christian S. Lentz, Mona Johannessen

## Abstract

The utilization of ATP within cells plays a fundamental role in cellular processes that are essential for the regulation of host-pathogen dynamics and the subsequent immune response. This study focuses on ATP-binding proteins to dissect the complex interplay between *Staphylococcus aureus* and human cells, particularly macrophages (THP-1) and keratinocytes (HaCaT), during an intracellular infection. A snapshot of the various protein activity and function is provided using a desthiobiotin-ATP probe, which targets ATP-interacting proteins. In *S. aureus*, we observe enrichment in pathways required for nutrient acquisition, biosynthesis and metabolism of amino acids and energy metabolism when located inside human cells. Additionally, the direct profiling of the protein activity revealed specific adaptations of *S. aureus* to the keratinocytes and macrophages. Mapping the differentially activated proteins to biochemical pathways in the human cells with intracellular bacteria revealed cell-type specific adaptations to bacterial challenges where THP-1 cells prioritized immune defenses, autophagic cell death, and inflammation. In contrast, HaCaT cells emphasized barrier integrity and immune activation. We also observe bacterial modulation of host processes and metabolic shifts. These findings offer valuable insights into the dynamics of *S. aureus*-host cell interactions, shedding light on modulating host immune responses to *S. aureus*, which could involve developing immunomodulatory therapies.

**Importance:** This study uses a chemoproteomics approach to target active ATP-interacting proteins and examines the dynamic proteomic interactions between *S. aureus* and human cell lines THP-1 and HaCaT. It uncovers the distinct responses of macrophages and keratinocytes during bacterial infection. *S. aureus* demonstrated a tailored response to the intracellular environment of each cell type and adaptation during exposure to professional and non-professional phagocytes. It also highlights strategies employed by *S. aureus* to persist within host cells. This study offers significant insights into the human cell response to *S. aureus* infection, illuminating the complex proteomic shifts that underlie the defense mechanisms of macrophages and keratinocytes. Notably, the study underscores the nuanced interplay between the host’s metabolic reprogramming and immune strategy, suggesting potential therapeutic targets for enhancing host defense and inhibiting bacterial survival. The findings enhance our understanding of host-pathogen interactions and can inform the development of targeted therapies against *S. aureus* infections.

Graphical Abstract

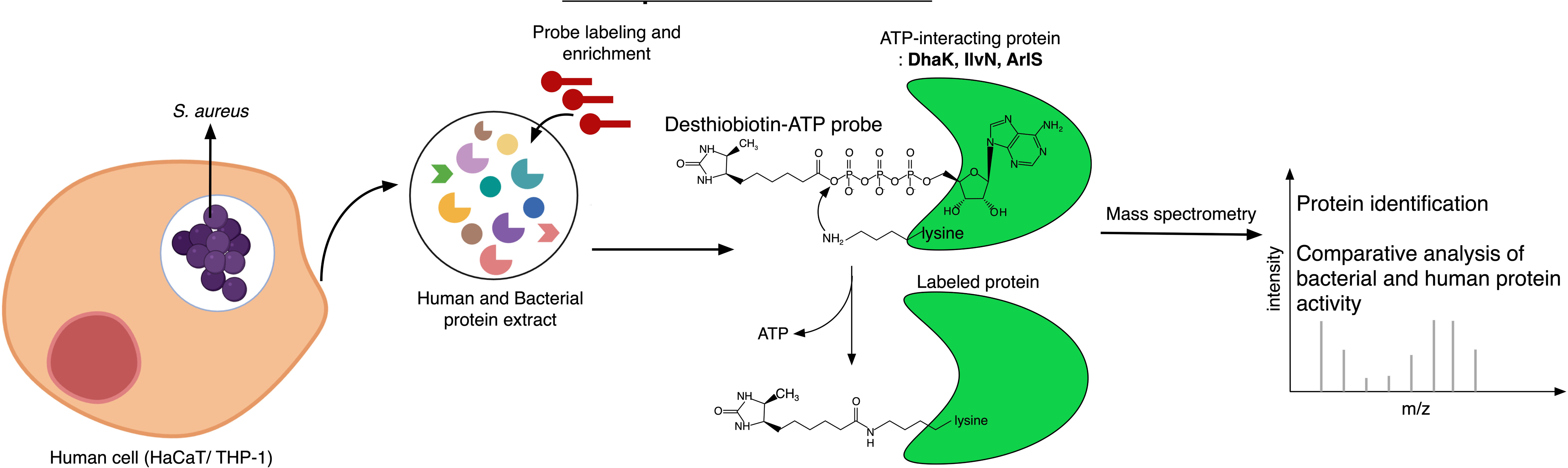

## Introduction

In the complex interface of host-pathogen dynamics, *S. aureus* infections across varied anatomical sites elicit a cascade of cellular responses*. S. aureus* causes conditions from minor skin irritations to life-threatening sepsis, altering its metabolism and virulence to thrive within host environments like macrophages and keratinocytes (1, 2). It counters host defenses by interrupting pathways like NF-kB and neutralizing ROS to protect itself from oxidative burst and antimicrobial peptides (AMPs). In keratinocytes, *S. aureus* alters surface proteins to evade recognition and secretes proteases that cleave AMPs, rendering them ineffective (3–6). *S. aureus* expertly exploits host mechanisms, manipulating ATP-binding proteins to alter inflammatory responses, potentially exacerbating immune reactions, causing tissue damage, and promoting its spread (7).

Keratinocytes, which guard the epidermal barrier, often serve as the first line of defense against microbial intruders. Meanwhile, professional macrophages patrol the systemic environment, carrying out specialized immunological functions, including phagocytosis and initiation of the adaptive immune response to eliminate pathogens (8). Bacterial infections trigger complex interactions between various immune cells and non-immune cells, utilizing distinct yet complementary mechanisms to recognize and eliminate pathogens. Professional phagocytes, notably macrophages, are pivotal in phagocytosis, antigen presentation, and directing immune responses whereas non-professional phagocytes, like keratinocytes, represent non-immune cells that are crucial in the defense against pathogens(9). Macrophages utilize pattern recognition receptors (PRRs) such as Toll-like receptors (TLRs) and Nod-like receptors (NLRs) to identify bacterial pathogen-associated molecular patterns (PAMPs) and initiate phagocytosis (10, 11). Inside these phagosomes, enzymes like NADPH oxidase generate reactive oxygen species (ROS), complemented by lysosomal acidification and proteases to eradicate bacteria (12, 13). Keratinocytes rely mainly on PRRs to produce AMPs and cytokines and chemokines to recruit immune cells. AMPs, particularly LL-37, can directly disrupt bacterial membranes (10, 14, 15). Upon PRR activation, both cell types trigger the NF-kB and inflammasome pathways, leading to an inflammatory response and the release of cytokines such as IL-1β and IL-18, highlighting their collaborative yet distinct contributions to immune defense (16, 17).

Macrophages detect pathogens and respond by reprogramming their metabolic pathways, known as the ‘immunometabolic shift.’ This shift facilitates various immune functions and pivots on the availability and utilization of ATP, a crucial energy storage molecule in living organisms(18, 19). ATP also plays a critical role in the signaling processes orchestrating these metabolic adjustments in both bacteria and host cells during infection. These metabolic switches are closely associated with specific ATP-interacting proteins that coordinate essential biochemical reactions (19, 20). Given their importance in cellular functions, these ATP-interacting proteins have been considered as key targets for antimicrobial development.

Unveiling the active fraction of ATP-interacting proteome during *S. aureus* infection in human cells through activity-based protein profiling (ABPP) can revolutionize our understanding of host-pathogen interactions. This chemoproteomic technique utilizes activity-based probes (ABPs) to capture and investigate the functional state of enzymes, transcending the limitations of transcriptomics and conventional proteomics by focusing on enzymatic activity rather than mere presence. The targeted enrichment involved in this chemoproteomic approach is also beneficial for identifying low-abundance proteins and functional characterization of hypothetical proteins (21). Nucleotide acyl phosphate probes are utilized for selective, covalent labeling and isolation of active kinases, ATPases and other ATP-binding proteins that are pivotal in human cellular processes, including both healthy and cancerous cells, as well as in microbial systems. This technique has facilitated the mapping of signal transduction pathways and provided insights into the regulatory mechanisms governing cell proliferation, apoptosis, and energy metabolism (22–25)

In this study, we utilized a desthiobiotin-ATP probe for the targeted profiling of ATP-interacting proteins in *S. aureus* and human cells during intracellular infection. Specifically, we used HaCaT cells, an immortalized human keratinocyte cell line, and THP-1 cells, a human monocytic cell line (THP1), to explore the biosynthetic and metabolic pathways adopted by bacteria during infection. Furthermore, we examined the differential immune responses elicited by these keratinocytes (non-professional phagocytes), and monocytes (professional phagocytes). This comparative approach to understanding the host-pathogen interplay during *S. aureus* infection provides profound insights into the pathogen’s pathogenesis and elucidates the complex molecular mechanisms that regulate these interactions.

## Materials and Methods

### Bacterial strains and growth conditions

The *S. aureus* strain used is the USA300_JE2, a derivative of a methicillin resistant *S. aureus* (MRSA) variant USA300_FPR3757. All strains used in this study are detailed in Table S1. *S. aureus* transposon mutants used in this study were provided by the Network on Antimicrobial Resistance in *Staphylococcus aureus* (NARSA) for distribution through BEI Resources, NIAID, NIH as part of the following reagent: Nebraska Transposon Mutant Library (NTML) Screening Array, NR-48501. For the infection of the human cell lines, the *S. aureus* strains USA300_JE2 was cultured in tryptic soy broth (TSB) at 37°C at 200 rpm. The overnight cultures were subcultured in fresh TSB and grown at 37°C at 200 rpm to an OD_600_ of 1.0. Bacterial pellets were washed twice with ice cold phosphate buffered saline (PBS) and resuspended in Roswell Park Memorial Institute (RPMI) or Dulbecco’s Modified Eagle’s medium (DMEM) depending on the experimental set up.

### Complementation of transposon mutants

The primers for cloning the genes for complementation are detailed in Table S2. The open reading frames (ORFs) of the genes were amplified from *S. aureus* USA300 JE2 genomic DNA by PCR, using Q5 Hot Start High-Fidelity DNA Polymerase (New England Biolabs). The resulting PCR products were then ligated into the EcoRI/KpnI sites of the pCM29-sgfp vector using the ClonExpress II One-Step Cloning Kit (Vazyme). These constructs were transformed into *E. coli* DH5α for screening and selection of the correct inserts by PCR. To improve the efficiency of transformation and overcome the restriction barrier, the recombinant plasmids were transformed into and extracted from *E. coli* IM01B. The resulting constructs were subsequently electroporated into the corresponding *S. aureus* transposon mutants and the transformants verified by PCR with plasmid-specific primers.

### Cell culture

HaCaT cells (Cytion) were cultured in DMEM with 10% fetal bovine serum (FBS) (Sigma) in T75 plates. THP-1 cells (TIB-202, ATCC) were cultured in RPMI 1640 media with 10% FBS (Sigma) in T75 plates. All cells were growth to ∼80% confluency in 37°C with 5% CO_2_. THP-1 monocyte differentiation was initiated by incubating the cells with 10 ng/mL phorbol 12-myristate 13-acetate (PMA; Sigma Aldrich) for 2 days at 37°C in a 5% CO_2_ environment. Following this 2-day differentiation period, the culture medium was aspirated, and the cells washed three times with PBS. Subsequently, the cells were further incubated for 1 day in fresh RPMI medium supplemented with 10% FBS.

### Infection of human cell lines for protein profiling

THP-1 macrophages and HaCaT cell lines were infected with the *S. aureus* USA300_JE2 at a multiplicity of infection (MOI) of 100. The human cells were incubated with bacteria for 1 hour at 37°C in the respective media, RPMI 1640 media for THP-1 and DMEM for HaCaT. The noninternalized bacteria were removed by washing the host cells 3 times with PBS, followed by incubation with fresh complete media of RPMI 1640 or DMEM supplemented with 100 μg/mL gentamicin for 1 hour at 37°C. The media was removed, and the cells washed three times with ice cold PBS and the cell culture plates placed on ice for protein extraction. As a negative control the THP-1 cells, HaCaT cells and bacteria were incubated separately in the DMEM and RPMI media.

### Profiling of human and *S. aureus* ATP-interacting proteins using desthiobiotin-ATP probe

Three sets of samples were prepared for each cell type to investigate the interaction with *S. aureus.* For the first set, THP-1 cells and HaCaT cells were infected with *S. aureus* in RPMI 1640 and DMEM media, respectively. The second set comprised THP-1 and HaCaT cells cultured alone in their respective growth media. The third set consisted of *S. aureus* cultured independently under identical conditions to those of the THP-1 and HaCaT cells (Fig. 1A). Following infection, cells were scraped of the plates into ice cold lysis buffer (25mM Tris HCl, 150mM NaCl, 1mM EDTA, 1% Nonidet P-40, 5% glycerol) with protease and phosphatase inhibitors. The cell suspension was transferred into 2 mL screw-cap tubes containing 100 μL of 0.1mm glass beads and lysed with bead-beating. The cell lysates were centrifuged for 20 min at 4°C and 14000 rpm. The supernatant (total lysate) was transferred to a new tube. The lysis buffer was exchanged for the reaction buffer (4M Urea+ lysis buffer) using the Zeba Spin Desalting Columns (ThermoFisher). Following buffer exchange the lysate protein concentration was measured using the Qubit protein assay kit and the protein concentration adjusted to 2 mg/ml using the reaction buffer. Each sample was mixed with 10 μL of 1M MgCl_2_ and incubated for 1 minute at room temperature. The AciveX^TM^ desthiobiotin-ATP probe (Thermo Fisher) was reconstituted in ultrapure water to 20 μM and 10 μL added to each sample followed by incubation for 10 minutes at room temperature. Following labeling 50 mL of 50% high-capacity streptavidin agarose resin slurry was added to each sample and incubated for 1 hour at room temperature with constant mixing on a rotator. The samples were centrifuged at 100x g for 1 minute to pellet the resin. The resins were washed 2 times with 500 mL of the reaction buffer and the bound protein eluted by adding 2X Laemmli reducing sample buffer and boiled for 5 minutes.

**FIG 1.**
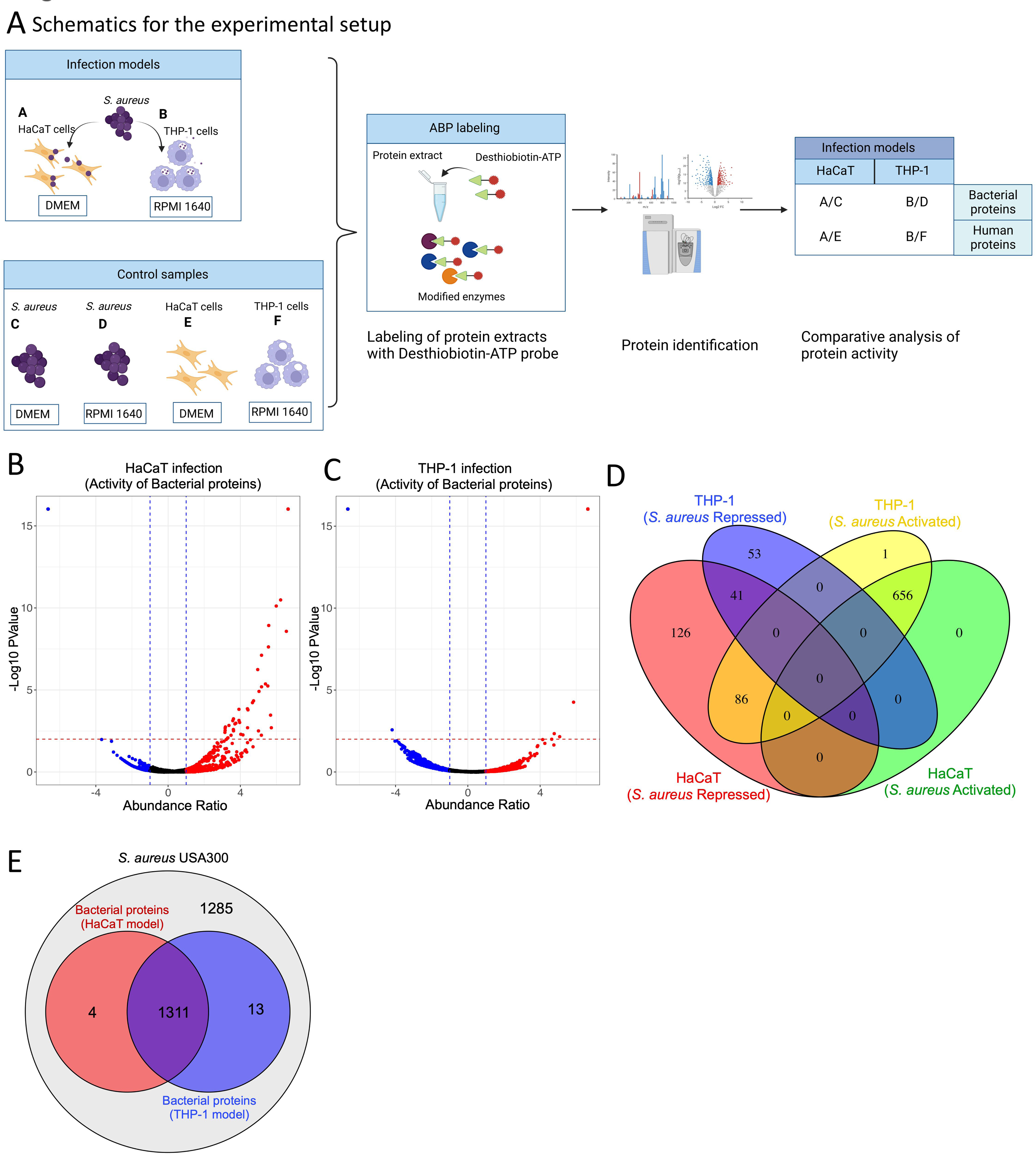
**A**) Schematic representation of the experimental setup including infection, ATP-interacting protein profiling and data analysis. The diagram gives an overview of the *S. aureus* infection of THP-1 and HaCaT cells and the control cultures of the human cells and bacteria grown in isolation. Probe labelling of the protein extracts from the samples was performed using the desthiobiotin-ATP probe and the modified proteins identified using mass-spectrometry. After infection, the differentially activated proteins in the *S. aureus* and host cells were identified by comparing them with the control samples, which are bacteria and human cell lines grown in isolation. **B**) and **C**) represent the activated and repressed proteins in the *S. aureus* following infection in the HaCaT and THP-1 cells and chemoproteomic enrichment, respectively. The abundance ratio is expressed as a log2 value of the relative activity of the proteins. **D**) Shows the differentially activated and repressed *S. aureus* proteins distribution following infection in the HaCaT and THP-1 cells. **E**) The general protein profile derived from the ATP probe in the *S. aureus* cell following infection in the THP-1 and HaCaT cells and their overlap with the total proteome in the *S. aureus* USA300 genome using the HMM profile derived from the UniProt database.

### Mass spectrometery and data analysis

Protein samples were initially reduced with dithiothreitol (DTT) and subsequently alkylated using iodoacetamide. Trypsin digestion was performed at a concentration of 6 ng/μl (Promega, #V511A). Post-digestion, peptides were purified with an Omix C18 tip (A57003100, Agilent), and the resulting eluates were dried via evaporation. For LC-MS analysis, the dried samples were reconstituted in 15 µl of 0.1% formic acid, and a quantity of 0.5 µg of peptides per sample was injected. The peptide mixtures, prepared in 0.1% formic acid, were loaded onto an EASY-nLC1200 system (Thermo Fisher Scientific) coupled with an EASY-Spray column (C18, 2µm, 100 Å, 50µm × 50 cm). A gradient of 5-80% acetonitrile in 0.1% formic acid was applied to fractionate the peptides over a 60-minute period at a flow rate of 300 nl/min. The fractionated peptides were then subjected to analysis on an Orbitrap Exploris 480 mass spectrometer (Thermo Scientific). Data acquisition was set to data-dependent mode, employing a Top40 method. For protein identification, the acquired data were searched against the *S. aureus* USA300_FPR3757 and human proteome databases using Proteome Discoverer 3.0 software. The search parameters included a peptide mass tolerance of 10 ppm and a fragment mass tolerance of 0.02 Da. A stringent FDR threshold of 5% was applied for peptide identification. To reinforce the reliability of the results, only peptides with at least two unique identifications were considered, and each sample was analyzed in triplicate to ensure reproducibility. The mass spectrometry proteomics data have been deposited to the ProteomeXchange Consortium via the PRIDE (26) partner repository with the dataset identifier PXD049003.

### Functional annotation and pathway analysis

To functionally annotate proteins differentially regulated in *S. aureus* and human cell lines (HaCaT and THP-1) post-infection, we integrated EggNOG for orthologous group classification, KEGG Pathway for metabolic and signaling roles, and gene ontology (GO) annotations. EggNOG provided insights into conserved domains and functions based on evolutionary history, while KEGG mapping highlighted their involvement in specific pathways. GO annotations were used to categorize proteins into biological processes (objectives the protein contributes to), molecular functions (biochemical activities), and cellular components (protein’s cellular location).

### Classification of ATP-interacting proteins in *S. aureus*

Protein sequences of *S. aureus* were retrieved from the UniProt database. A comprehensive set of known ATP-binding motifs was compiled from literature and existing protein databases. These motifs were used to construct a Hidden Markov Model (HMM) profile using the HMMER software suite to detect ATP-binding domains. The complete proteome of *S. aureus* USA300 FPR3757 genome and the total proteome from the HaCaT and THP-1 infection models was screened against the constructed HMM profile. This was aimed to identify potential ATP-interacting proteins by matching the amino acid sequences of the proteins to the ATP-binding HMM profile.

### Galleria infection assay

Larvae of the *Galleria mellonella* were obtained from Reptilutstyr AS (Norway). For infection experiments, *S. aureus* strains were grown overnight in TSB, washed twice in PBS, and suspended in PBS to a final concentration of 1.0 × 10^5^ CFU/ml. Larvae of approximately equal weight were inoculated with 20 µl of this bacterial suspension, resulting in an infection dose of 1×10^3^ CFU/larva. Bacteria were injected into the hemocoel of the larvae between the last pair of legs using a 30G syringe microapplicator (0.30 mm (30G) × 8 mm, BD Micro-Fine demi). As a control, larvae were mock-inoculated with 20 µl of a PBS solution. Survival of the larvae was observed for every 3 hours at 37°C. Larvae were regarded dead when they were not moving upon repeated physical stimulation.

### Time-course human cell line infection assay

For the time-course infection assay, THP-1 and HaCaT cells were inoculated with the *S. aureus* strains at a MOI of 10. The THP-1 and HaCaT cells were cultured for 1 hour with the bacteria in RPMI 1640 and DMEM, respectively without antibiotics. To eliminate non-internalized bacteria, the cells were washed thrice with PBS and then incubated in fresh complete media containing 50 μg/mL gentamicin. At designated time points, the media was discarded, the cells washed with PBS, lysed with 0.01% Triton X-100 in PBS, and the lysates were serially diluted and plated on Mueller Hinton agar. After 24 hours of incubation at 37°C, the bacterial colonies were counted.

### Time-course human cell line cytotoxicity assay

THP-1 and HaCaT cells were infected with the *S. aureus* strains at a MOI of 10 for 1 hour within RPMI 1640 and DMEM, respectively. Subsequently, cells were washed thrice with PBS to eliminate non-internalized bacteria. They were cultured in fresh RPMI 1640 or DMEM media, each supplemented with 50 μg/mL gentamicin, throughout the time-course assay. Post-incubation, the media were aspirated, cells were washed thrice with PBS, and the MTT (3-(4,5-dimethylthiazol-2-yl)-2,5-diphenyltetrazolium bromide) reagent diluted in the respective cell culture media, added to the cells, followed by incubation at 37°C for 2 hours. The wells were washed twice with PBS, and DMSO was added to solubilize the formazan crystals. Alongside these samples, control assays with uninfected cells were conducted to ensure assay validity (Fig. S1). The absorbance at 520 nm was measured using a BioTek Synergy H1 plate reader.

### Statistical Analysis

Normality and homogeneity of variance were assessed using Shapiro-Wilk and Levene’s tests, respectively. For data that was both normally distributed with homogenous variances, a one-way ANOVA followed by Dunnett’s post hoc test was used for comparisons against the wildtype (WT). Data that were not normally distributed were analyzed using the Kruskal-Wallis test followed by Dunn’s post hoc test for multiple comparisons.

## RESULT AND DISCUSSION

### Profiling of ATP-interacting proteins in *S. aureus*

Employing a desthiobiotin-ATP probe facilitated the targeted investigation of proteins exhibiting altered activity profiles during *S. aureus* infection in human keratinocyte (HaCaT) and monocyte (THP-1) cell lines. Proteomic samples from these cells, both independently cultured and infected with *S. aureus*, were labelled with the probe for enrichment and subsequent analysis. By comparing the proteomic data from the infected state to that of cells grown under controlled media conditions, we delineated the activity profile of both bacterial and human proteins (from HaCaT and THP-1 cells) that were activated or repressed during infection (Fig. 1A). To enhance the selectivity of our chemoproteomic analysis using the desthiobiotin-ATP probe, which exhibits potential cross-reactivity with various metabolic enzymes and chaperones, we implemented a stringent exclusion criterion. Proteins exhibiting consistency with the predetermined threshold cut-offs in all biological replicates were retained. This selective approach yielded a set of proteins with consistent activity across the experimental conditions, facilitating accurate comparative analysis across our experimental and control datasets. The combined datasets (Tables S3) show the bacterial proteins resulting from the verification of activity-dependent protein interactions in our study.

*S. aureus* exhibited distinct protein activation profiles when infecting the HaCaT and THP-1 cells (Fig. 1B-D). While a significant number of proteins (656) were activated in both HaCaT and THP-1 cells, only 41 proteins were commonly repressed across both cell models. Interestingly, 86 proteins activated in THP-1 cells corresponded with those repressed in HaCaT cells. In contrast, the THP-1 model showed repression of 53 bacterial proteins, whereas in the HaCaT cell, 126 *S. aureus* proteins were repressed, indicating unique bacterial responses to different host cell environments (Fig. 1D). Furthermore, approximately half of the proteins from *S. aureus* USA300 were identified in the infection models using the ATP probe (Fig. 1E).

To categorize the differentially activated protein sets identified using the ATP probe, we developed an HMM using data from the Uniprot database, focusing on well-known bacterial protein domains that interact with ATP or nucleotides. Subsequently, we organized the proteins from each model into their respective functional or structural categories. Among the identified proteins, a substantial group comprised proteins with nucleotide-binding domains, with the majority showing no overlap with proteins possessing well-defined ATP-interacting domains (Fig. S2). The overlapping classification of ATP-interacting proteins can be attributed to certain proteins containing multiple domains. Furthermore, enriched proteins that did not match any of the defined ATP-or nucleotide-binding domains were classified as “Unassigned,” which mainly encompassed hypothetical proteins, proteins with poorly annotated domains and metabolic enzymes. The molecular basis for the interaction of these proteins with the ATP-probe is unclear and it cannot be ruled out that the enrichment of some of these targets is the result of non-specific interactions, in which case the observed differences between different biological conditions represent differences in ‘abundanc’ rather than in ‘activit’ (Fig. S2). It is also possible that some proteins may have been enriched indirectly based on their interaction with other ATP-probe labelled enzymes within molecular complexes. However, nucleotide acyl phosphate probes have been shown previously to reveal novel ATP-binding sites where ATP might act as allosteric regulators of enzyme function (27). Allosteric control of enzyme activity through ATP is common for metabolic enzymes such as phosphofructokinases PfkA (28, 29) and might also be potential novel function for unassigned proteins in our dataset. The specificity of these interactions may be addressed in follow-up studies e.g. using a competition assay with ATP and other nucleotides (27, 30). The unassigned proteins provide valuable insights into uncharacterized proteins with ATP-interacting functions, potentially shedding light on their roles in metabolism, stress responses, and host-pathogen interactions.

### Functional classification of bacterial enzymes

We employed a comprehensive approach combining enrichment analysis, GO, and cluster of ortholog (COG) analysis for a broader perspective on protein functions by considering evolutionary relationships, metabolic pathways, and molecular interactions. Based on the COG analysis, the enrichment of proteins in categories including amino acid transport and metabolism (E), energy production and conversion (C), transcription (K), translation and ribosomal biogenesis (J), replication, recombination, and repair (L), cell wall/membrane biogenesis (M), and carbohydrate transport and metabolism (G) (Fig. 2A) The comparative analysis using the COG categories of the activated and repressed bacterial proteins from the HaCaT and THP-1 infections shows that specific functional categories are more affected by the bacterial presence in one cell type over the other, which indicates a targeted bacterial strategy for adapting to specific host cellular environments. Specifically, it was observed that in the THP-1 model, there is a higher number of activated bacterial proteins in most categories than in the HaCaT model. On the other hand, in the HaCaT model, there are more repressed proteins as compared to the THP-1 model (Fig 2A), suggesting cell-type-specific interactions and bacterial response strategies and could also indicate a strategic bacterial adaptation to the intracellular environment of that specific host cell type. For example, the category related to transcription (K) and translation (J) shows a substantial number of repressed proteins in the HaCaT model (Fig 2A). This repression could reflect *S. aureus* adaptive response by downregulating proteins involved in transcription and translation to conserve energy and resources or avoiding the host’s immune detection mechanisms.

**FIG 2.**
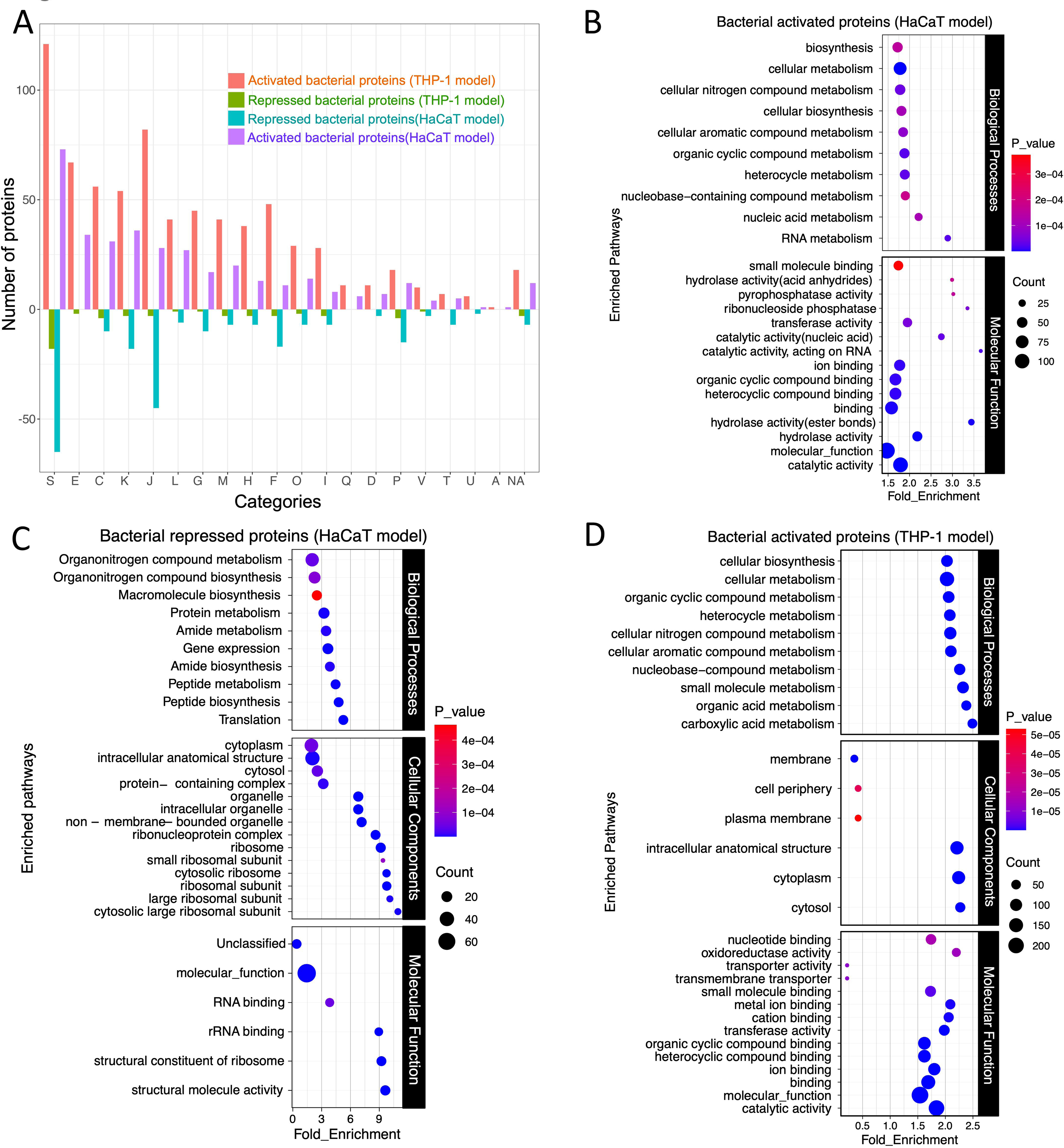
A) COG analysis of the activated and repressed *S. aureus* proteins following infection in the THP-1 and HaCaT cells and chemoproteomic enrichment. The length of the Up and Down bars represents the number of activated and repressed proteins, respectively. The bar chart illustrates the differential activation of *S. aureus* proteins categorized by COG during infection in the THP-1 and HaCaT cells. Each bar represents the degree to which proteins within a specific COG category are either activated or repressed. The x-axis labels, denoted by single-letter codes, correspond to distinct functional COG categories. The GO analysis of the **B)** bacterial activated and **C)** repressed proteins following infection in the HaCaT cells and **D)** the bacterial activated proteins following infection in the THP-1 cells.

Additional GO analysis of the differentially active bacterial proteins from HaCaT and THP-1 infection models revealed enrichment in metabolic pathways utilizing various biomolecules such as carboxylic and organic acids and organic substance biosynthesis (Fig. 2B and D). The distribution of molecular functions in *S. aureus* proteins after infection in HaCaT and THP-1 indicates their diverse roles, including catalytic activities, transport functions, and binding activities, suggesting involvement in enzymatic reactions, molecule transport, and interactions with ions and small molecules (Fig. 2B-D). During infection in the HaCaT cells, most enriched bacterial processes included proteins involved in catalytic activities, RNA-related functions, phosphatase activity, and acid anhydride hydrolase activity (Fig 2B). In contrast, in the THP-1 infection, proteins involved in molecular functions such as transporter activity, oxidoreductase activity, nucleotide binding, and metal ion binding were prominent (Fig. 2B-D). These distinctions suggest that in the HaCaT model, the bacteria activate proteins related to enzymatic and RNA-related processes, while in the THP-1 infection, most activated proteins function in transport, redox reactions, and nucleotide interactions.

During infection in the HaCaT cells, the repression of bacterial proteins associated with translation, peptide, and protein metabolism when they are located intracellularly (Fig. 2C) suggests a downregulation of the bacterial protein synthesis machinery, which could conserve energy or divert resources toward other processes necessary for survival or immune evasion. This aligns with the activation of pathways involved in RNA and nucleic acid metabolism (Fig. 2B), indicating an increased focus on genetic regulation and adaptation by the bacterium. Additionally, the downregulation of bacterial molecular pathways like translation, peptide biosynthesis, gene expression and macromolecule and organonitrogen compound biosynthesis during *S. aureus* infection in HaCaT cells indicates a shift in bacterial metabolic priorities (Fig. 2C).

### Bacterial energy metabolic pathways during intracellular location

Investigation into bacterial protein-mediated metabolic signatures, prompted by the enrichment of energy production and conversion pathways during infections, revealed their critical role in catabolic pathways typical of intracellular pathogens. These pathways include fatty acid degradation, the tricarboxylic acid cycle, glycolysis, pyruvate metabolism, and formate production (Fig. 3A).

**FIG 3.**
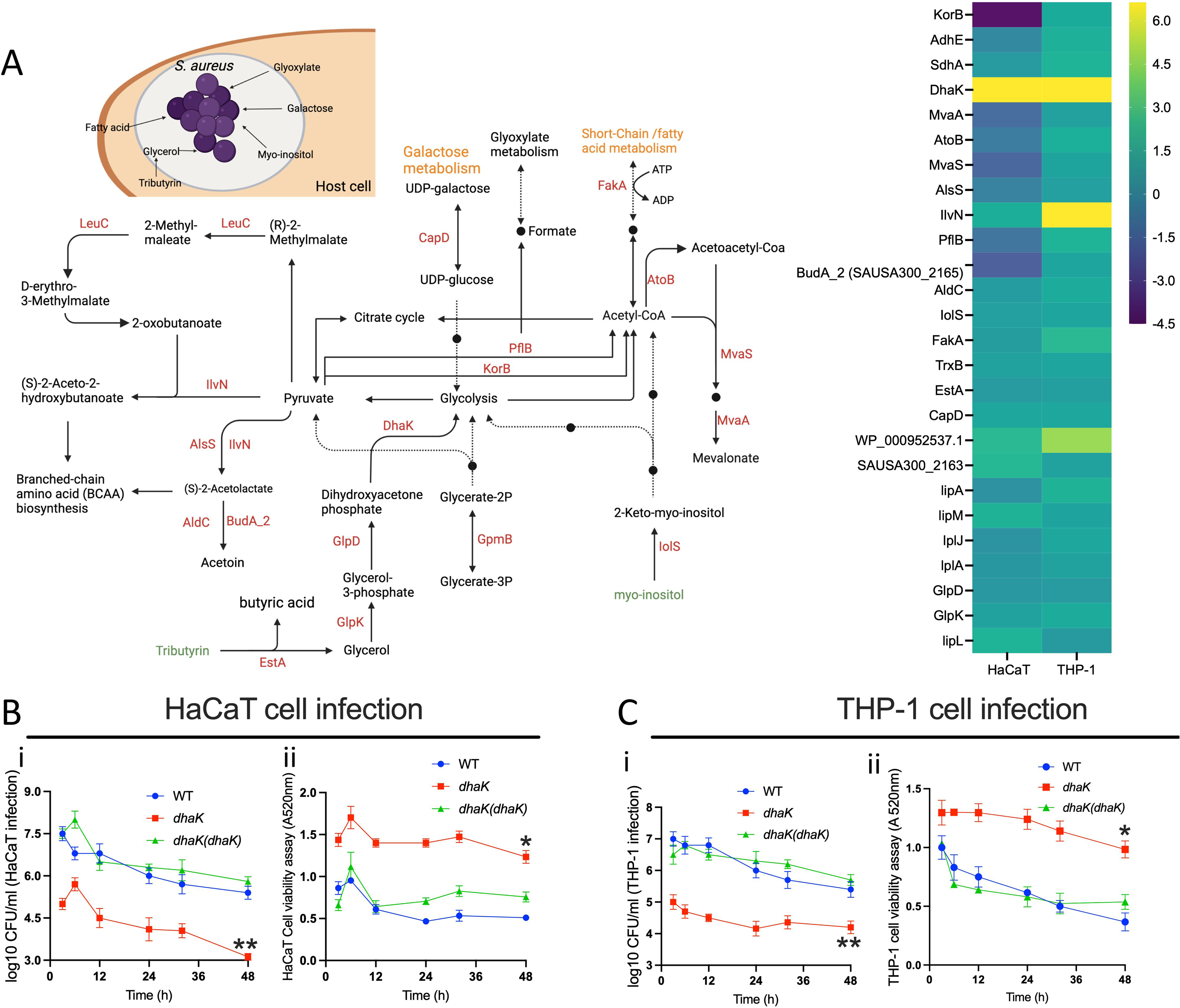
**A**) The metabolic map and heatmap show *S. aureus* proteins involved in energy metabolism and carbon source utilization after infection in HaCaT and THP-1 cells. The heatmap shows the relative activity of the bacterial proteins. The heatmap scale shows the log2 value for the protein activity. The map renders the pathways derived from the KEGG and EggNog databases. **B)** and **C)** The time-course HaCaT and THP-1 infection and cell viability assays following infection with the *S. aureus* strains; the WT, transposon mutant of *dhaK* and its in-trans complemented strain (*dhaK(dhaK)*). The data represents the mean ± SD of three independent experiments. A P-value > 0.05 is considered significant. (P<0.01 is indicated as *, and P< 0.001 is indicated as **).

While some intracellular bacteria, such as *Listeria monocytogenes* and *Mycobacterium tuberculosis*, may prefer glycerol and fatty acids as carbon sources, respectively (31), our observations in *S. aureus* infecting both HaCaT and THP-1 cells indicate significant activation of DhaK, a dihydroxyacetone kinase involved in glycerol metabolism (32)(Fig 3A-B). We also observed increased enrichment of EstA/FphF an esterase that we recently identified in another chemoproteomic study using serine hydrolase-reactive fluorophosphonate probes (33, 34), but this esterase does not possess domains likely to directly interact with ATP. Increased activity of EstA and DhaK, associated with the utilization of tributyrin via glycerol, suggests that glycerol may serve as a significant carbon source for metabolic processes during intracellular infection. In support of this, in *L. monocytogenes,* a mutant strain lacking DhaK, which is defective in glycerol and dihydroxyacetone (DHA) utilization, has been shown to exhibit a reduced intracellular replication rate (35, 36). Using our *S. aureus* strain background, we confirmed that a DhaK transposon mutant displayed a reduced ability to infect/replicate in both THP-1 and HaCaT cells (Fig. 3B (i) and 3C (i)). Moreover, this mutation correlates with a decrease in cytotoxicity in these cell lines (Fig 3B (ii) and 3C (ii)). Complementation with DhaK-expressing plasmid rescued the phenotypes in both cell lines (Fig. 3B and C).

Proteins like CapD and IolS, involved in galactose and myo-inositol catabolism respectively, were activated in both HaCaT and THP-1 infections (Fig. 3A), highlighting the bacterium’s ability to catabolize host-derived molecules to fuel its metabolic pathways (37, 38).

In both HaCaT and THP-1 cells, the activation of proteins for lipoic acid biosynthesis (LipA, LipL, LipM) and scavenging (LplA, LplJ) was observed, with higher activity noted in THP-1 cells (Fig. 3A and B). This suggests that *S. aureus* relies on both its innate biosynthesis and external scavenging mechanisms for lipoic acid during infection, which is key for its defense against reactive oxygen and nitrogen species from macrophages, aiding in its evasion of immune clearance (39, 40). Additionally, in both HaCaT and THP-1 cell infection, bacterial protein FakA involved in fatty acid and acetate metabolism (41) showed increased activity, indicating a preference for fatty acid metabolism (Fig. 3A and B).

The enzyme PflB, which converts pyruvate to acetyl-CoA and formate under anaerobic conditions was uniquely active in the THP-1 model (Fig. 3A and B), while KorB (SAUSA300_1183), also involved in pyruvate to acetyl-CoA conversion, was activated in THP-1 (42, 43). This dual activation of PflB and KorB for pyruvate metabolism could indicate a metabolic flexibility or redundancy in *S. aureus* approach to energy production, allowing it to adapt to the intracellular environment of the macrophages.

Furthermore, the enzymes AtoB, MvaS, and MvaA (Fig. 3A) play a crucial role in the mevalonate biosynthesis pathway and were found to be activated during infection in THP-1 cells but repressed during HaCaT infection (Fig. 3B). In *S. aureus*, mevalonate is essential for the primary metabolism of bacteria, and it is the only route for the biosynthesis of isoprenoids, which are crucial for cell wall formation, respiratory energy generation, and oxidative stress protection (44, 45).

### Bacterial amino acid metabolism and transport during intracellular location

Intracellular bacteria like *S. aureus* can de novo synthesize or scavenge amino acids for metabolic energy, cellular integrity, and biosynthesis of vital compounds, including nucleotides and cell wall components (31, 46, 47).

During HaCaT cell infection, we noted an elevated activity of the *S. aureus* amino acid transport system BrnQ (SAUSA300_1300), higher than during THP-1 cell infections (Fig. 4A and B). BrnQ is essential for the uptake of branched-chain amino acids, including leucine, isoleucine, and valine (48). To evaluate this, we infected HaCaT and THP-1 cells with WT *S. aureus,* a transposon mutant of *brnQ* and its complemented strain. The *brnQ* mutation markedly diminished *S. aureus* ability to infect both THP-1 and HaCaT cells over prolonged periods. This mutation simultaneously reduced the cytotoxic effects on the cells. The complementation of *brnQ* in these mutants restored their infective proficiency and cytotoxicity to wild-type levels (Fig. 4C (i and ii) and 4D (i and ii)), confirming the effect of the mutation.

**FIG 4.**
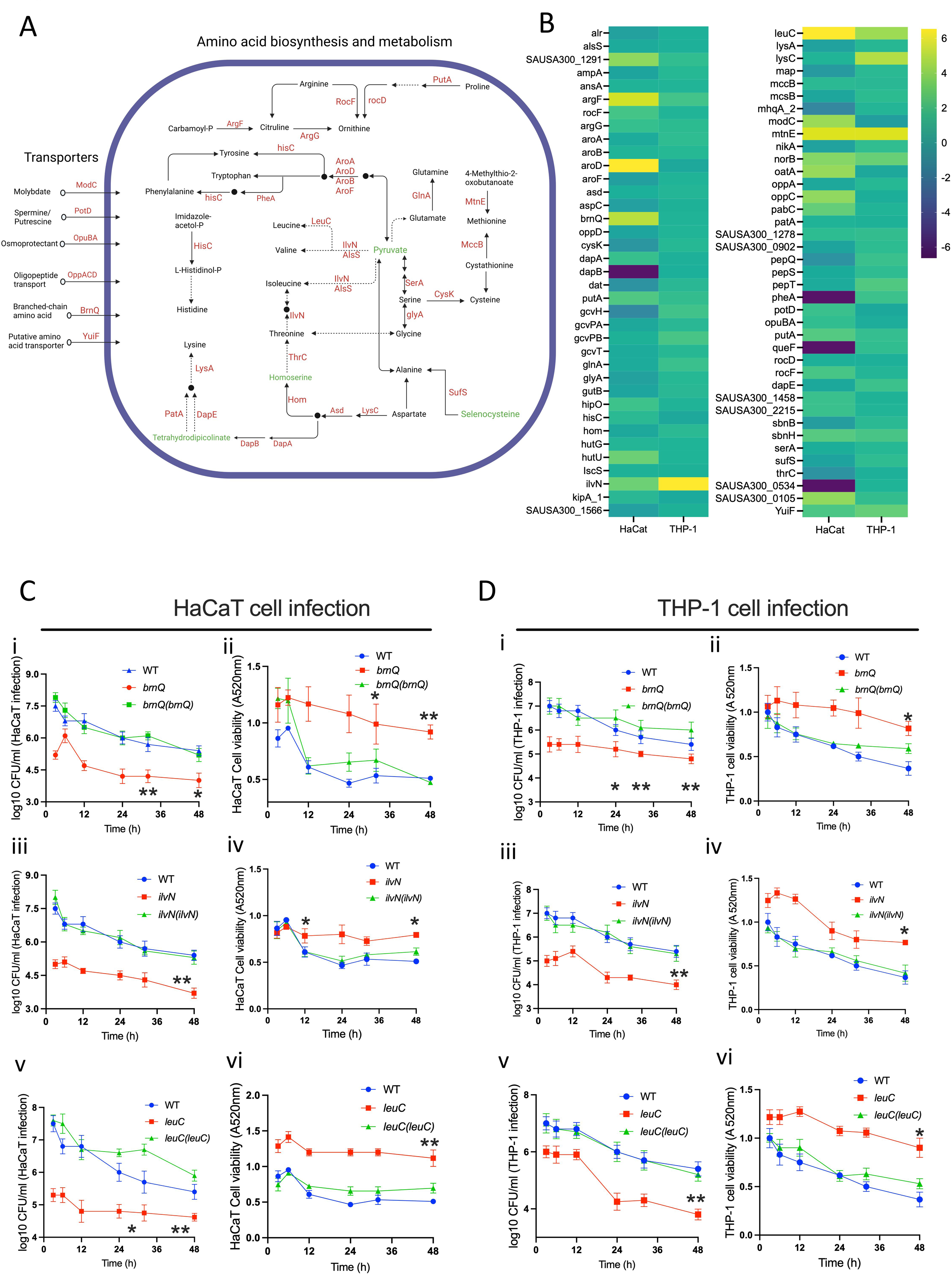
**A**) A diagrammatic rendering of the amino acid synthesis and transport pathway involving bacterial proteins activated and repressed following infection in the THP-1 and HaCaT cells. **B)** The heatmap shows the relative activity (log2 value) of bacterial proteins involved in the amino acid synthesis, metabolism, and transport pathway. **C)** and **D)** The time-course HaCaT and THP-1 infection and cell viability assay following infection with the WT *S. aureus* strains, transposon mutants of *brnQ*, *IlvN* and *leuC* and their corresponding complemented strains *brnQ(brnQ)*, *ilvN(ilvN)* and *leuC(leuC)*. The data represents the mean ± SD of three independent experiments. A P-value > 0.05 is considered significant. P<0.01 is indicated as *, and P< 0.001 is indicated as **.

Simultaneously, IlvN and LeuC, involved in the biosynthesis of branched-chain fatty acids, were activated during infection in both HaCaT and THP-1 cells (Fig. 4A and B), indicating a dual strategy of biosynthesis and scavenging by the bacteria in response to intracellular location. The transposon mutants of both *ilvN* and *leuC* display reduced infection capabilities and reduced cytotoxicity in both HaCaT and THP-1 cell lines during prolonged infection (Fig 4C (iii-vi) and 4D (iii-vi)). IlvN, in conjunction with other activated proteins like AlsS, BudA, and AldC, plays a vital role in acetoin production (Fig. 3A and S3). This compound acts as a storage form of surplus carbon and energy and is generated during glucose fermentation. Additionally, it serves as an external electron acceptor during anaerobic conditions (49, 50).

AroD, a key enzyme in the shikimate pathway involved in the biosynthesis of aromatic amino acids (phenylalanine, tyrosine, and tryptophan), was highly enriched during HaCaT cell line infection compared to in THP-1 cells (Fig. 4A and B). In some other bacteria, the AroD-catalyzed product 3-dehydroshikimate, combined with citrate and spermidine, contributes to synthesizing the siderophore petrobactin which is however not reported for *S. aureus* (51). Yet, the high activation of PotD, a protein involved in spermidine transport, in bacteria during HaCaT infection may indicate a convergence of these metabolic pathways also in *S. aureus* (Fig. 4A and B). During both HaCaT and THP-1 infections, the AroA and AroB, which are responsible for downstream chorismate production (52), showed lower levels of activity. On the other hand, PheA, which is involved in phenylalanine biosynthesis, was activated during THP-1 infection but repressed during HaCaT infection. The MtnE, and MccB involved in methionine salvage pathway (53) exhibited similar activation levels during infection in both HaCaT and THP-1 cell lines (Fig. 4A and B).

The proteins GcvH, GcvT, GcvPA, and GcvPB, which are involved in glycine metabolism to lipoyllysine, showed increased activity during THP-1 infection compared to the HaCaT cell infection (Fig. 4B and S3). Lipoyllysine is essential for intermediary metabolism, including the tricarboxylic acid cycle, and is required for the breakdown of glucose and the generation of acetyl-CoA(54). Given the observed activation of proteins related to lipoic acid biosynthesis and salvage pathways in *S. aureus* during its infection of human cells (Fig 3B and S3), it is likely that lipoic acid-dependent post-translational modifications, particularly those involving the glycine cleavage system play an essential role in the bacterial survival strategy within the host cells.

Additionally, HutU, which is involved in histidine metabolism and L-glutamate, aspartate, and alanine biosynthesis (Fig. 4B and S3), was activated in the HaCaT model and the RocF/Arg, which participates in arginine degradation to urea (55), was activated in the HaCaT infection model (Fig. 4B and S3).

### Bacterial virulence and stress response during infection of kerationocytes and macrophages

Central to bacterial infections is the expression of specific proteins that play pivotal roles in virulence, direct damage to host tissues, orchestration of nutrient acquisition in nutrient-limited conditions, and the subversion of the host’s immune defenses. Here SbnE, involved in staphyloferrin B production(56), and the major facilitator superfamily transporter NorB, showed significant activation in both infection models (Fig. S3). In the HaCaT infection model *S. aureus* proteins such as RadA (a DNA repair protein) and SAUSA300_2299 (a multidrug resistance protein A transporter) were highly activated. PflB involved in response to NO stress (57) was activated during THP-1 infection, which aligns with macrophage reactive nitrogen species (RNS) production (Fig. S3). Furthermore, proteins related to oxidative stress response, like BshB (involved in Bacilithiol biosynthesis), LepA (influencing vacuolar modeling for host immune evasion), and RocF (involved in putrescine and urea production necessary for reactive nitrogen species detoxification) were identified. Additionally, proteins such as SAUSA300_2140, IsdB, and PerR, associated with iron-siderophore biosynthesis, were also found to be activated (Fig. S3). Beyond the well-known pathways enriched based on GO, KEGG, and COG categories, various characterized and uncharacterized bacterial proteins mediating stress response, virulence and survival during infection showed differential activity in both models (Fig. S4).

### Bacterial two component systems activated during infection in HaCaT and THP-1 cells

The bacterial two-component systems (TCS) and other kinases sense the environment, enabling bacteria to adapt accordingly. Profiling the bacterial proteins allowed us to identify a set of histidine kinases including some belonging to the TCS as well as serine-threonine kinases in *S. aureus*.

In the HaCaT infection model, the TCS sensor kinases ArlS, NreB, SrrB, and WalK exhibited increased activity, whereas the GraS was repressed (Fig. 5A). Furthermore, during infection in THP-1 cells, the histidine kinases ArlS, SrrB, WalK, were upregulated (Fig. 5B). Notably, the accessory component YycI, which forms a complex WalKR-YycHI to regulate virulence determinants and stress response pathways in *S. aureus*, showed increased activity in both infection models (Fig. S3). Using the *Galleria mellonella* infection model, we confirmed that transposon mutants deficient in the histidine kinase or response regulators of the SrrAB and ArlSR, showed increased survival of infected larvae compared to WT strains (Fig. 5C). To further investigate the role of the TCS in host infection, we infected THP-1 and HaCaT) cells with transposon mutants and complements of the TCSs, SrrAB and ArlSR. During the prolonged infection, we observed that the mutants of both sensor kinases and response regulators exhibited reduced infection capacity and cytotoxicity in both cell lines. However, this attenuated virulence was restored to the WT level in their respective complemented strains (Fig. 5D and E). The WalKR, an essential TCS, was not included due to the unavailability of corresponding mutants.

**FIG 5.**
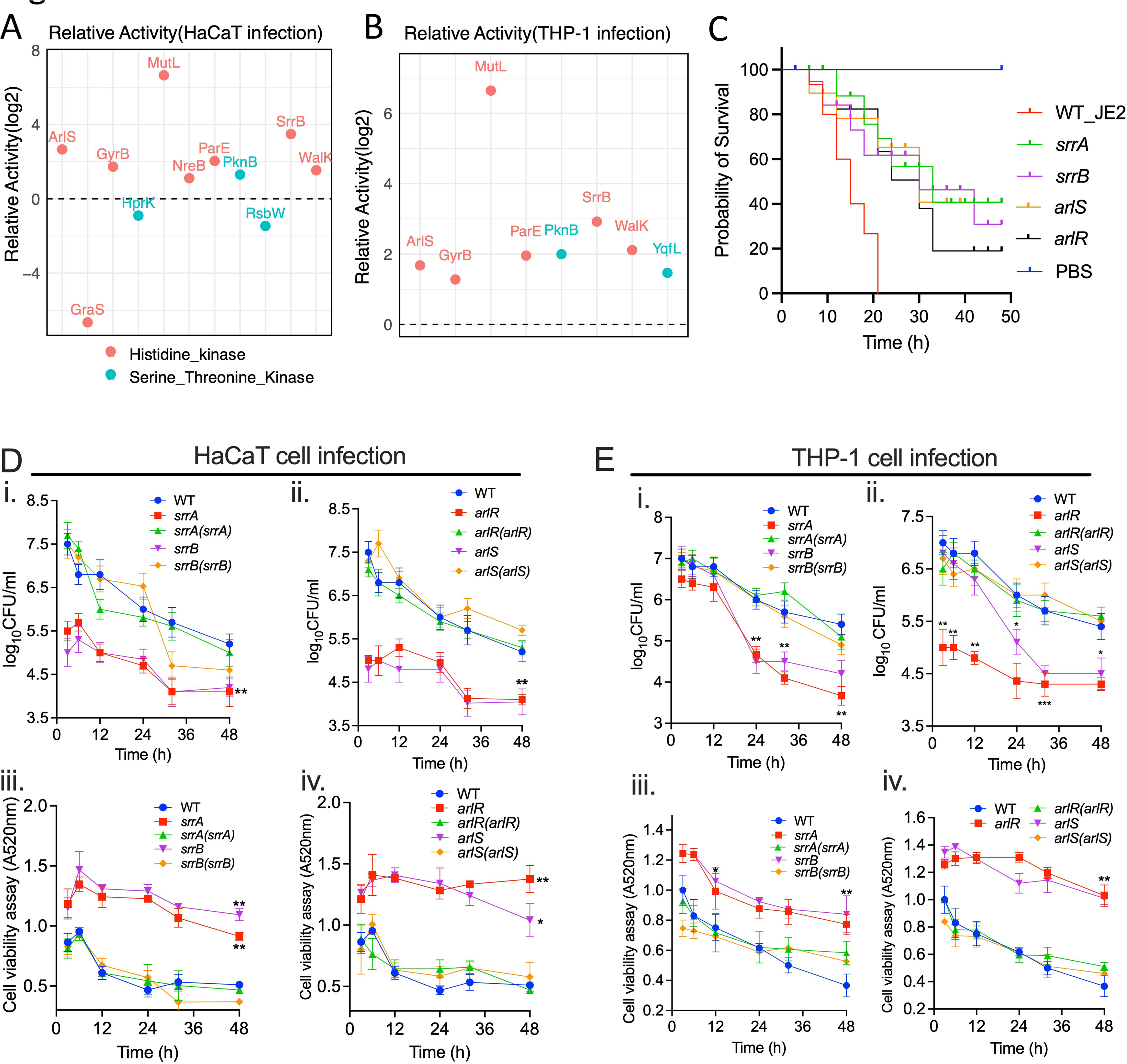
Relative activity of *S. aureus* histidine kinases and serine/threonine kinases following infection in **A)** HaCaT cells and **B)** THP-1 cells. **C)** The *Galleria mellonella* survival following infection by the wildtype *S. aureus* USA300_JE2 and the transposon mutants of the TCSs SrrAB and ArlSR. PBS was used as a control. **D**) and **E**) The time-course HaCaT and THP-1 infection and cell viability assay following infection with the WT *S. aureus* strains, transposon mutants of *srrA*, *srrB, arlR,* and *arlS*, and their corresponding complemented strains *srrA(srrA), srrB(srrB)*, *arlS(arlR),* and *arlR(arlR)*. The data represents the mean ± SD of three independent experiments. A P-value > 0.05 is considered significant. P<0.01 is indicated as *, and P< 0.001 is indicated as **.

In both infection models, *S. aureus* serine-threonine kinase PknB was active, whereas YqfL activation was specific to THP-1 cell infections. Conversely, HprK, a serine kinase and RsbW, a serine-threonine-protein kinase were slightly suppressed during HaCaT cell infection (Fig. 5A and B). These kinases are vital for regulating cellular processes; for example, HprK regulates carbon metabolism, PknB is tied to cell growth, division, and cellular quiescence, and RsbW is associated with the stress response (58–61). Also, the GyrB, ParE and MutL were all activated in bacteria during infection in both cell lines (Fig 5A and B). ParE and GyrB are both involved in the manipulation of DNA supercoiling and are part of topoisomerase complexes, whereas MutL is involved in the repair of DNA mismatches (62, 63). All three are essential for the maintenance and regulation of the bacterial genome integrity.

### Distribution and functional categorization of human proteins in mammalian cells infected with ***S. aureus***

After *S. aureus* infection, THP-1 and HaCaT cells displayed cell-specific responses in their profile of activated and repressed proteins (Fig. 6A). Functional categorization revealed that a significant number of proteins are associated with categories such as K (transcription), T (signal transduction mechanisms), J (translation, ribosomal structure, and biogenesis), C (energy production and conversion), and O (post-translational modification, protein turnover, chaperones) (Fig. 6B) reflecting the host’s effort to control the infection by regulating immune responses, metabolic adaptation, and ensuring protein functionality. Interestingly, there were minimal overlaps in the activated and repressed proteins between the two cell lines (Fig. 6A and B), signifying that while there are some common elements in the response to *S. aureus* infection, each cell type also employs unique strategies that are likely tailored to their specific roles in the immune system. The combined datasets (Tables S4) show the THP-1 and HaCaT proteins resulting from the verification of activity-dependent protein interactions in our study.

**FIG 6.**
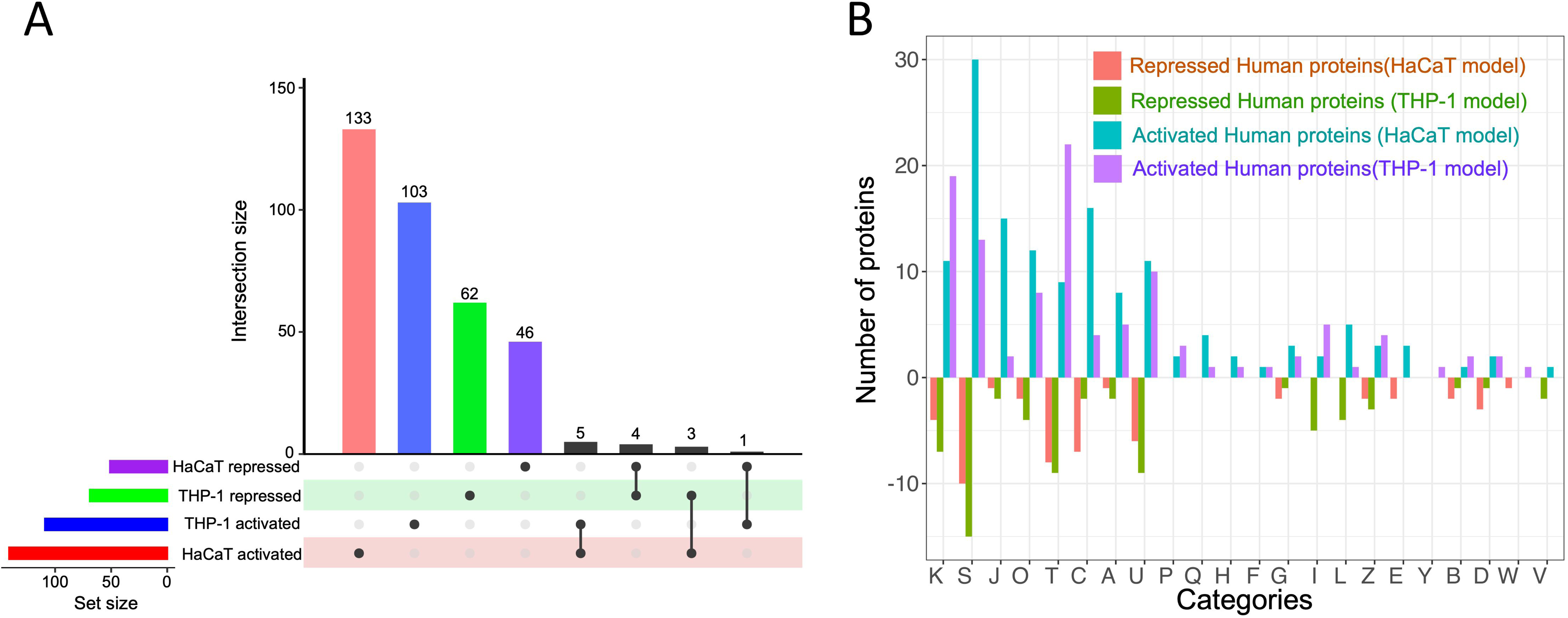
**A**) Distribution of activated and repressed human proteins from HaCaT and THP-1 cells following *S. aureus* infection. **B**) This bar chart illustrates the differential activation of human proteins categorized by Clusters of Orthologous Groups (COG) in THP-1 and HaCaT cells in response to *S. aureus* infection. Each bar represents the degree to which proteins within a specific COG category are either activated or repressed. The x-axis labels, denoted by single-letter codes, correspond to distinct functional COG categories.

### Functional profiling of HaCaT proteins in response to intracellular *S. aureus* infection

The gene ontology and biochemical pathway analysis of the differentially activated proteins from the HaCaT cell following *S. aureus* infection profoundly impacted cellular energy metabolism and signaling. Many of the activated proteins in the HaCaT cells were involved in mitochondrial processes, including mitochondrial translation, gene expression, respiratory chain complex assembly, and electron transport (Fig. 7A). This suggests that *S. aureus* infection in HaCaT cells may lead to an increase in ATP production or indicate mitochondrial stress or damage induced by the bacteria. Also, the enrichment in pathways related to oxidative phosphorylation, thermogenesis, and electron transport to ubiquinone suggests an alteration in energy metabolism and potential oxidative stress in the infected cells (64–66)(Fig 7A and B). The activation of the chemical carcinogenesis-reactive oxygen species (ROS) pathway in HaCaT cells during *S. aureus* infection involved the CYBA which is involved in ROS production and MGST3 which protects against oxidative stress (67–69) and IKBKG directly involved in the activation of NF-κB pathway (Fig 7B)(16, 70). Various components of the mitochondrial electron chain transport components and NADH dehydrogenase complex (NDUFS7, NDUFA8, NDUFA3, NDUFB1, NDUFC2, NDUFS5) (Fig. 7A and B), indicate that the cells are potentially trying to eliminate the bacteria through oxidative stress and the energy demands required for immune response (71–74).

**FIG 7.**
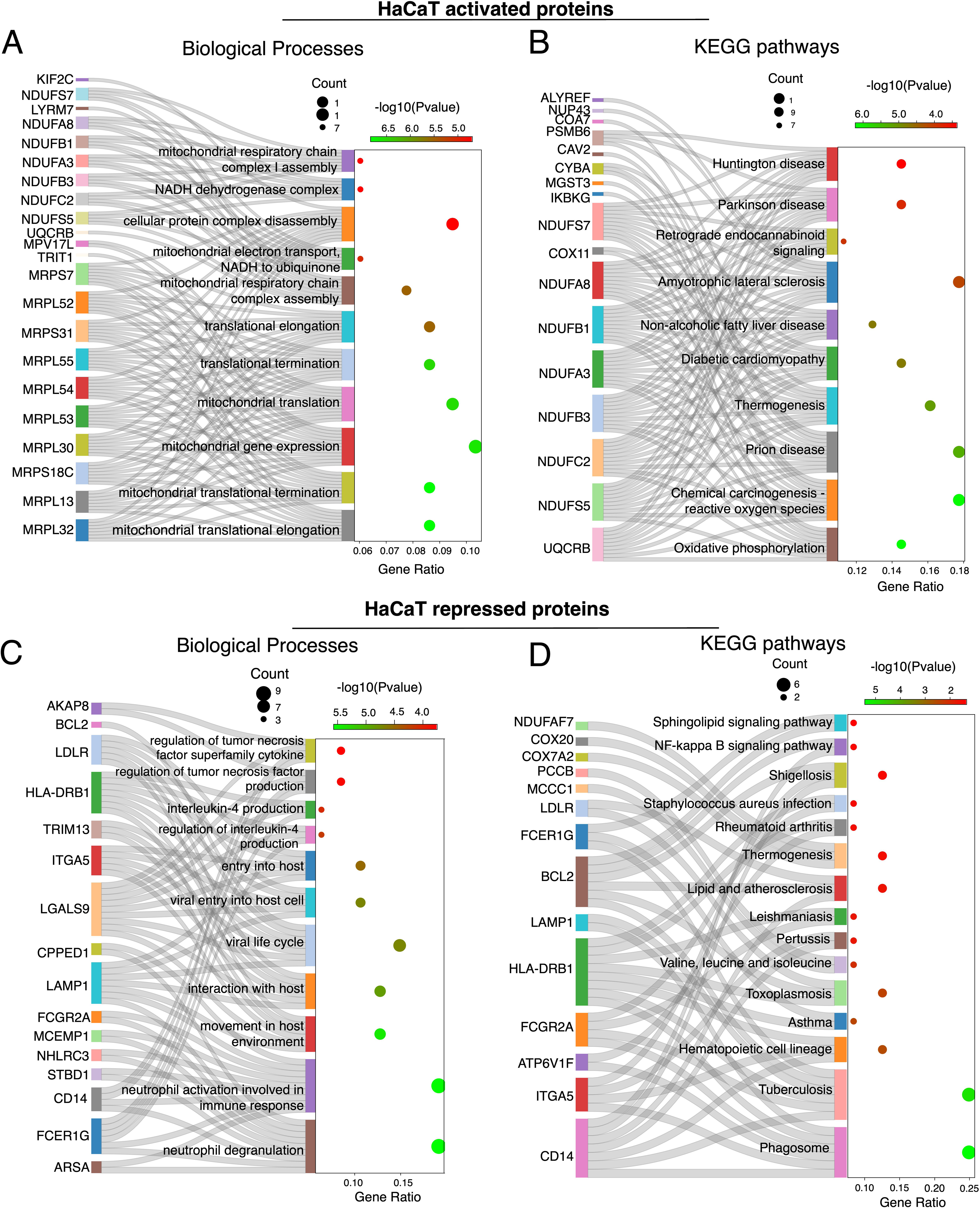
Sankey-dot plot showing the flow of activated proteins to their respective **A**) biological processes and **B**) KEGG pathway enrichment in HaCaT cells following *S. aureus* infection. **C)** Sankey-dot plots depicting the repressed proteins and their corresponding biological processes and **D**) KEGG pathway enrichment in HaCaT cells following *S. aureus* infection. The dot size indicates gene count and color intensity represents the-log10(P-value).

Annotations for cellular components and molecular functions showed a strong influence on mitochondrial aspects, including the mitochondrial inner membrane, mitochondrial ribosome, mitochondrial respiratory chain complexes, and NADH dehydrogenase complex I (Fig. S5A and B), indicating that the infection has a significant impact on energy metabolism and ATP demand. Pathways related to oxidoreductase activity, particularly those affecting NAD(P)H and enzymes like glutathione peroxidase, suggest potential redox imbalance and oxidative stress (68, 69). Additionally, pathways for binding activities such as transferrin receptor and long-chain fatty acid binding indicate shifts in cellular transport and metal ion homeostasis (Fig S5A and B). In general, the activated protein in the HaCaT cells indicate that *S. aureus* may either disrupt mitochondrial energy production or provoke the cell to enhance mitochondrial function as a protective measure.

Conversely, proteins repressed in HaCaT cells post-infection were significantly enriched in pathways related to immune response alterations, intracellular trafficking, and energy metabolism (Fig. 7C and D). This included downregulation of neutrophil-related pathways (neutrophil activation involved in immune response and neutrophil degranulation) and cytokines like interleukin-4, TNF and the repression of proteins involved in the phagosome pathway (Fig. 7C and D) (75, 76). Additionally, the repression in pathways annotated for shigellosis, tuberculosis, and toxoplasmosis and even *S. aureus* infections (involving BCL2, FCGR2A, HLA-DRB1, FCER1G, CD14, LAMP1 that have functions in apoptosis, phagocytosis, and inflammatory response) (Fig. 7D), may reflect a deviation of the immune system induced by *S. aureus* (75–79).

Interestingly, the ATP-interacting protein profiles of host and bacteria give evidence for their metabolic interdependencies that might be indicative of the competition of resources. Whereas *S. aureus* upregulates BrnQ branched chain amino acid (BCAA) transport systems, likely to sequester BCAAs for its growth (Fig. 4A and B), the repression of MCCC1 and PCCB (Fig. 7D), that are crucial for BCAA catabolism, suggest that HaCaT cells try to conserve BCAAs for immune function and tissue repair (80, 81).

Finally, the repression of cellular components related to endocytic vesicles, secretory granules, and lysosomes could impact antigen processing and pathogen defense, while the downregulation of binding activities such as immunoglobulin and ubiquitin-binding may affect immune recognition and protein degradation processes (Fig. S5C and D).

### Functional profiling of THP-1 proteins in response to intracellular *S. aureus* infection

THP-1 cells responded to *S. aureus* infection through changes in lipid metabolism, DNA integrity, and immune response. The enriched pathways included those related to cholesterol and sterol transport, suggesting active modulation of lipid metabolism during infection (Fig. 8A and B). Cholesterol and sphingolipid metabolism are essential for membrane structure, signaling pathways, and apoptosis (82). Additionally, WRNIP1, TP53BP1, CLU, SMCHD1, CBX8, and RINT1 were chemoproteomically enriched in *S. aureus* infected THP1 cells (Fig. 7E). These proteins are involved in responding to DNA damage, chromatin regulation, and double-strand break repair, indicate potential DNA damage due to bacterial invasion or modulation by the host to limit bacterial replication (83–87). Furthermore, proteins like SPG7, MFN1, VPS18, C2CD5, EEA1, USP8, DNAJC13, GOLGA2, and SEC24B were enriched (Fig. 8A), and these are critical for organelle fusion and vesicle organization, play vital roles in cellular processes such as autophagy, apoptosis, lysosomal degradation of pathogens, antigen presentation, and cytokine release, all essential for the immune system’s defense against bacterial infection (88, 89).

**Fig 8.**
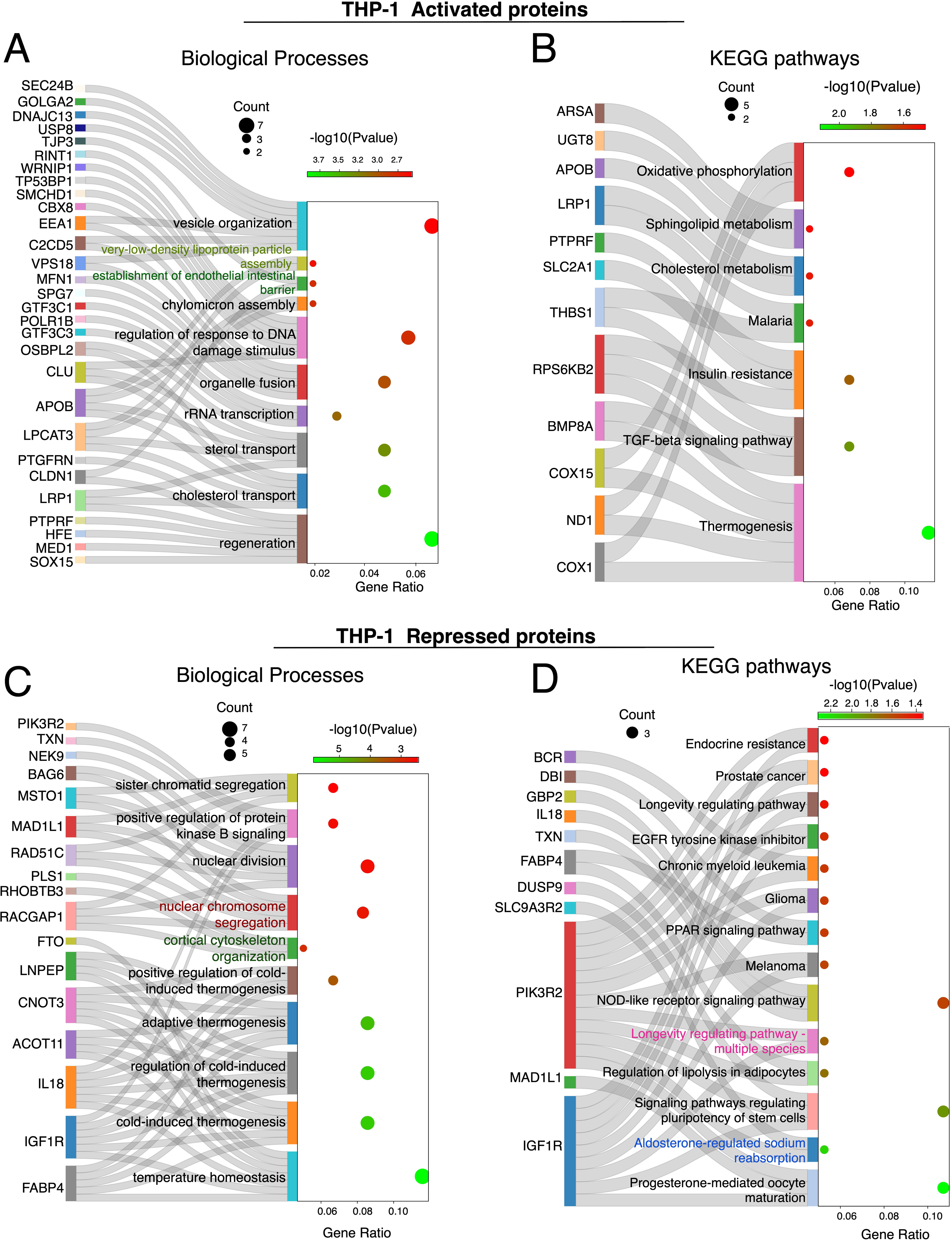
Sankey-dot plot illustrating the activated proteins and their related **A**) biological processes and **B**) KEGG pathway enrichment in THP-1 cells following *S. aureus* infection. **C)** Sankey-dot plots of repressed proteins and their biological processes and **D**) KEGG pathway in THP-1 cells following *S. aureus* infection. The dot size indicates gene count and color intensity represents the-log10(P-value).

Further analysis of activated THP-1 proteins revealed their roles in lipid transport and metabolism involving proteins such as CLU, OSBPL2 and APOB(90–92), cell membrane dynamics (early endosome, coated vesicle, vesicle tethering complex, vesicle lumen and desmosome), oxidoreductase activities, and heme-copper terminal oxidase activity (Fig. S6A and B), which are associated with mitochondrial function, the electron transport chain, cellular energy metabolism, and responses to oxidative stress (93, 94). Regarding energy production and metabolism, COX1, ND1, COX15 play roles in energy metabolism and mitochondrial function (95, 96)(Fig. S6A and B).

The repressed proteins in THP-1 cells during *S. aureus* infection might indicate disruptions in cellular processes including metabolism, cell division, signaling, and intracellular trafficking. While cold-induced or adaptive thermogenesis typically involves converting energy into heat through proteins like ACOT11, FABP4, IL18, IGF1R, CNOT3, and FTO (97–101) (Fig. 8C), this process bypasses ATP synthase, releasing energy as heat rather than ATP (102, 103). However, due to the energy demands of the THP-1 cell response to bacterial infection, processes like thermogenesis, which consume energy without ATP production, are suppressed (102). Thus, during infection, the cell prioritizes energy pathways that support ATP production, crucial for powering immune responses. This metabolic shift underscores the cell’s adaptive ability to ensure survival during infection-induced stress.

The KEGG pathways analysis of the THP-1 repressed proteins revealed enrichment in pathways such as progesterone-mediated oocyte maturation, endocrine resistance, melanoma, glioma, chronic myeloid leukemia, prostate cancer, EGFR tyrosine kinase inhibitor resistance, and longevity regulating pathways, primarily involving proteins PIK3R2 and IGFIR linked to the PI3K/Akt signaling pathway (Fig. 8D)(104). Furthermore, proteins related to vesicles and endosomes, critical for cellular trafficking, material exchange, and DNA replication, were suppressed during bacterial infections (Fig. S6D). This may reflect a bacterial strategy to inhibit cell proliferation and potentially manipulate cell death to their advantage (105). Additionally, proteins facilitating molecular functions, such as GTPase and small GTPase binding, essential for intracellular trafficking, signaling, and cytoskeletal organization, were also repressed (Fig. S6C).

### Immune and metabolic shifts in THP-1 and HaCaT cells during bacterial challenge

Protein mapping in host cells exposed to *S. aureus* reveals distinct, cell-type specific immune adaptations. THP-1 cells activated proteins associated with immune defense, autophagy, and inflammation. In contrast, HaCaT cells activated proteins that support cellular barrier integrity and trigger immune responses, reflecting tailored strategies to bacterial invasion (Fig. 9).

**FIG 9.**
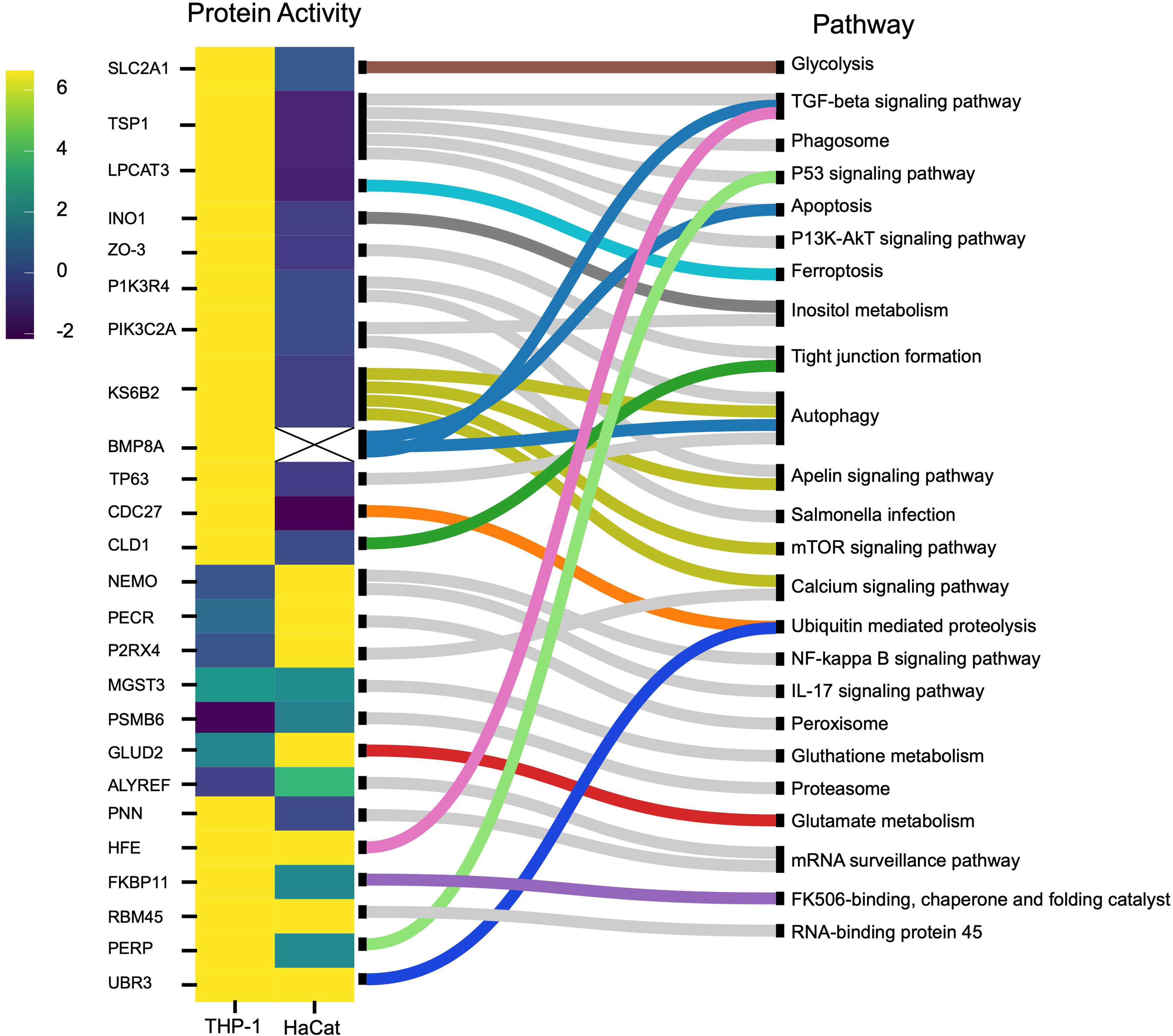
**A**) List of human immune response pathways. Heatmap and pathway mapping of differentially activated human proteins from the HaCaT and THP-1 cells following *S. aureus* infection. The heatmap scale shows the log2 value for the protein activity.

Upon examining the link between protein activity and pathways in THP-1 cells, we observed a prioritization of immune defense mechanisms, including enhanced pathogen clearance and immune activation, autophagic cell death, and inflammatory responses during pathogen exposure. Proteins such as TSP1, LPCAT3, BMP8A, TP63, PIK3R4, and KS6B2 play a crucial role in autophagy, apoptosis, and ferroptosis facilitating pathogen clearance, remove compromised cells, and limit the spread of infection (106–111). Also, following exposure to pathogens, PERP, an apoptosis-inducing protein (112, 113), showed higher activation levels in THP-1 cells than in HaCaT cells (Fig. 9). Furthermore, proteins like INO1 and PIK3C2A for inositol synthesis, essential in cellular signaling, growth, proliferation, differentiation, survival, and intracellular trafficking, were activated in THP-1 cells upon infection (114, 115). Notably, myo-inositol also plays a role in depolarizing macrophages, enhancing phagocytosis (116), and inducing autophagy independently of mTOR (mammalian target of rapamycin) signaling pathway (117)

Exposure to pathogens has been found to change the metabolic pathway of host cells. In this case, the repression of SLC2A1, a glucose transporter, in HaCaT cells and its activation in THP-1 cells reflect their metabolic needs for immune response (Fig. 9). SLC2A1 is involved in glycolysis by supporting glucose uptake and promoting anaerobic metabolism via the hypoxia-inducing factor (HIF)-signaling pathway (118, 119). Therefore, by limiting glucose uptake, HaCaT cells can restrict the availability of glucose, which is a major carbon source for *S. aureus* metabolism. This innate defense strategy is an example of “nutritional immunity” (120). Additionally, PECR, involved in fatty acid oxidation was repressed in THP-1 cells, but activated in HaCaT cells (Fig. 9). This leads to a shift in the metabolic reprogramming of immune cells during infection from fatty acid metabolism to glycolysis, known as the Warburg effect (121, 122). Many intracellular pathogens have evolved to take advantage of the Warburg metabolism of the host cell or to push the host cell into a state of increased glycolysis (121, 122). The Warburg metabolism, mediated by the mTOR/HIF-1alpha pathway, is crucial for producing IL-22 and cytokines during intracellular infections (98, 99).

Regarding nutritional immunity, host cells often limit iron availability to inhibit bacterial proliferation. Both LPCAT3 (activated only in THP-1 cells) and HFE (activated in both HaCaT and THP-1 cells) (Fig. 9) proteins are involved in iron regulation mechanisms via the iron-dependent programmed cell death (ferroptosis) pathway (124–126).

HaCaT cells engage in mechanisms that mitigate excessive inflammation and cellular damage, preserving cellular integrity and the skin’s protective barrier. The upregulation of NEMO, PECR, and P2RX4, implicated in NF-κB, IL-17, peroxisome, and calcium signaling pathways (Fig. 8), demonstrates the HaCaT cells’ strategic response to *S. aureus*. This includes bolstering the immune response via the NF-κB pathway and enhancing antimicrobial and inflammatory actions through IL-17 signaling. NEMO, through NF-κB, is essential for keratinocyte survival, proliferation, and migration in response to various stressors such as cytokines, bacterial and viral products (127–129). In contrast to the THP-1 cells, the proteins PIK3R4, KS6B2, BMP8A, TSP1, LPCAT3, and TP63 were suppressed in HaCaT cells. These proteins are involved in autophagy, *Salmonella* infection, phagosome, ferroptosis, apoptosis, and the P53 signaling pathway (116). PECR, a peroxisome reductase, is involved in fatty acid synthesis (130), vital in forming the epidermal permeability barrier, and keratinocytes function as building blocks and energy generation, storage, and metabolism (131, 132). The activation of PECR in HaCaT cells (Fig. 9) may suggest an increased synthesis of fatty acids to maintain keratinocytes layer disrupted by the infection. PNN located in epithelial cell desmosomes, regulates cell integrity and adhesion (133). Host factors such as NEAT-1, associated with PNN, triggers macrophage inflammation (134, 135). Silencing NEAT-1 by ALYREF repression reduces ATP and mitochondrial energy production (136). Additionally, PNN overexpression creates a stable adhesive state, impeding cell transition and affecting epithelial tissue structure and function (137). Hence, suppressing PNN can help HaCaT cells maintain the immune barrier and facilitate post-bacterial challenge recovery.

The differential activities of the proteins in both HaCaT and THP-1 cells could result from the bacteria’s modulation of host immune processes as an immune evasion mechanism. For instance, disrupting the tight junction due to repression of CLD1 and ZO-3 in HaCaT cells (Fig. 9) can have immediate implications for cell integrity and could represent the success of bacteria in compromising barriers, and facilitating invasion and progression of infection. *S. aureus* is known to disrupt tight junctions to facilitate invasion and spread (138, 139). Also, various bacteria have been reported to subvert host defenses to promoting inflammation, depending on the bacterial needs and the phase of infection by targeting the NF-κB pathway (124)

## Conclusion

This study delves into the interplay between ATP-interacting proteins in *S. aureus* and human cells, specifically THP-1 macrophages and HaCaT keratinocytes, during an infection. By employing chemical probes designed to target specific proteins selectively, our research unveils distinctive patterns of protein activation within *S. aureus* and human cells highlighting the strategies and adaptations during host-pathogen interactions. A graphical summary of the pathways and proteins activated during the *S. aureus*-human cell interaction is illustrated in Fig. S7.

These findings pinpoint potential targets for therapeutic interventions and contribute to a more comprehensive understanding of the dynamics that unfold during *S. aureus* infections. This knowledge serves as the foundation for developing interventions to disrupt bacterial virulence or fortify the host’s defense mechanisms, thereby aiding in the ongoing battle against antibiotic resistance. Moreover, the insights gained into *S. aureus* metabolic adjustments during infection hold promise for the creation of novel antimicrobials that target these pathways. Additionally, our research unveils strategies to modulate host immune responses, potentially leading to the development of immunomodulatory therapies that enhance the host’s defense mechanisms or mitigate damage caused by inflammation. The findings of this study open diverse avenues for future research, for a deeper exploration of molecular-level interactions between hosts and pathogens.

## Acknowledgements

This work was funded through a grant from the Centre for New Antibacterial Strategies (CANS) to K.H., M.H. and C.S.L., and a CANS start-up grant to C.S.L. Mass spectrometry-based proteomic analyses were performed by UiT Proteomics and Metabolomics Core Facility (PRiME). This facility is a member of the National Network of Advanced Proteomics Infrastructure (NAPI), which is funded by the Research Council of Norway INFRASTRUKTUR-program (project number: 295910).

Nebraska transposon mutant library (NTML) was provided by the Network on Antimicrobial Resistance in *Staphylococcus aureus* (NARSA) for distribution through BEI Resources, NIAID, NIH: NTML Screening Array, NR-48501.

